# Regional glia-related metabolite levels are higher at older age and scale negatively with visuo-spatial working memory: A cross-sectional proton MR spectroscopy study at 7 tesla on normal cognitive ageing

**DOI:** 10.1101/864496

**Authors:** Anna Lind, Carl-Johan Boraxbekk, Esben Thade Petersen, Olaf Bjarne Paulson, Hartwig Roman Siebner, Anouk Marsman

**Author notes:** Author contributions: A.L., A.M., E.P., C.J. and H.S. designed the research, A.L., A.M and E.P. established and piloted the MR-spectroscopic measurements and A.L. carried out the experiments. A.L., A.M., C.J., and H.S. analysed the data. All authors participated in the interpretation of the data, A.L. wrote the paper draft, all authors reviewed and edited the paper. Corresponding author: Anouk Marsman, Danish Research Centre for Magnetic Resonance (DRCMR), Copenhagen University Hospital Hvidovre, Section 714, Kettegård Allé 30, 2650 Hvidovre, Denmark.

## Abstract

Proton MR spectroscopy (^1^H-MRS) has been used to assess regional neurochemical brain changes during normal ageing, but results have varied. Exploiting the increased sensitivity at ultra-high field strength, we performed ^1^H-MRS in 60 healthy volunteers to asses age-related differences in neuron- and glia-related metabolite levels and their relation to cognitive ageing. Gender was balanced and participants were assigned to a younger, middle and older group according to their age, ranging from 18 to 79 years. They underwent ^1^H-MRS of anterior cingulate cortex (ACC), dorsolateral prefrontal cortex (DLPFC), hippocampus and thalamus and performed a visuo-spatial working memory task outside the scanner. Multivariate ANCOVA revealed a significant overall effect of age group on metabolite levels in all brain regions of interest. Considering specific metabolites, elevated levels of glia-related metabolites in older age groups were observed in all brain regions. For the neuron-related metabolites, differences among age groups were only observed in two regions. We found higher levels of total N-acetyl aspartate in hippocampus for the older than the middle group, and lower levels of glutamate in DLPFC in the older than the younger group. Working memory performance correlated negatively with levels of glia-related metabolites for total creatine and total choline in ACC and myo-inositol in hippocampus and thalamus, but not with neuron-related metabolites. Together, the results show that age effects on glia-related metabolites scale with cognitive ageing in multiple brain regions. We conclude that ^1^H-MRS-derived glia-related markers may be of value in longitudinal studies of brain ageing.

**Significance statement:** Neurochemical ageing is an integral component of age-related cognitive decline. Proton MR spectroscopy (^1^H-MRS) studies of *in vivo* neurochemical changes across the lifespan have, however, yielded inconclusive results. Due to the higher spectral resolution and sensitivity, 7 tesla ^1^H-MRS has potential to improve consistency of the results. Here we apply this technique to demonstrate that glia-related metabolites are elevated in older age in anterior cingulate cortex, hippocampus and thalamus with a novel link to cognitive ageing of visuo-spatial working memory. This study is the first to investigate normal ageing in these brain regions using 7 tesla ^1^H-MRS and the results indicate that glia-related metabolites could be of value in longitudinal studies of cognitive ageing.

## Introduction

Describing normal human ageing is vital for understanding what distinguishes both successful and pathological ageing. Normal ageing involves cognitive decline related to widespread neurochemical alterations (Driscoll et al., 2003). These alterations have been studied non-invasively and *in vivo* using proton MR spectroscopy (^1^H-MRS), however, the results have been highly variable (for review see: (Haga et al., 2009; Cichocka and Bereś, 2018; Cleeland et al., 2019)).

The most commonly investigated brain metabolites in ^1^H-MRS ageing studies are myo-inositol (mIns), total creatine (tCr), total choline (tCho), total N-acetylaspartate (tNAA) and glutamate (Glu) (Cichocka and Bereś, 2018). Though results from ^1^H-MRS ageing studies have varied, the most consistent results indicate that mIns, tCr and tCho levels increase with age whereas tNAA and Glu levels decrease (Cleeland et al., 2019). As mIns, tCr and tCho have higher levels in glial cells than neurons, the increase potentially reflects gliosis and neuroinflammation (Glanville et al., 1989; Urenjak et al., 1993; Chang et al., 2013). tNAA and Glu levels are commonly regarded to reflect neuronal health and function, suggesting that a decrease in their levels is linked to compromised neuronal integrity (Clark, 1998; Demougeot et al., 2001; Rae, 2014). Due to the variation in results, this ageing pattern is, however, not conclusively established.

One reason for the previous variation in results could be that the neurochemical changes during ageing are highly region dependent (Eylers et al., 2016). In some regions, such as the thalamus, age-related changes are rarely observed (Gruber et al., 2008; Yang et al., 2015; Eylers et al., 2016) whereas in PFC and hippocampus age effects are frequently observed though the specific metabolites and the direction of the effect varies (Schubert et al., 2004; Chiu et al., 2014; Ding et al., 2016; Sporn et al., 2019). The proposed pattern of age-related changes may, thus, primarily involve some brain regions more sensitive to ageing.

There are a number of additional potential reasons for the variability of the results, such as the included age-spans, sample sizes and scan parameters (Cichocka and Bereś, 2018). Further, many studies quantify metabolite levels by calculating the ratio to tCr. As tCr is indicated to change with age, this complicates interpretation of the results (Cleeland et al., 2019). Moreover, the distribution of grey matter (GM), white matter (WM) and cerebrospinal fluid (CSF) within a given ^1^H-MRS voxel depends on each individual’s anatomy and the extent to which this is corrected for may influence the results. Lastly, the use of lower field strengths (1.5-4T) impact the sensitivity, specificity and spectral resolution compared with ^1^H-MRS at higher field strengths such as 7T (Tkac et al., 2001; Terpstra et al., 2016). Developments in ^1^H-MRS hardware and best practice could, thus, increase the consistency and interpretability of the results.

The overall aim of this study was to investigate regional brain metabolite differences across three age groups of normal individuals using 7T ^1^H-MRS and to relate the differences in metabolite levels to cognitive ageing. The primary hypothesis was that the metabolite levels would differ across age groups in a region-dependent manner. More specifically, it was predicted that in the regions proposed to be more ageing-sensitive, namely anterior cingulate cortex (ACC), dorsolateral prefrontal cortex (DLPFC) and hippocampus, older age groups would show higher levels of mIns, tCr and tCho and lower levels of tNAA and Glu. In thalamus, no differences were expected. The secondary hypothesis was that ageing-sensitive metabolites would be associated with cognitive ageing. Visuo-spatial working memory (vsWM) was chosen to exemplify cognitive ageing based on the literature showing how this function is sensitive to ageing and depends on the regions of interest in this study (Owen et al., 1996; Brockmole and Logie, 2013; Goldstone et al., 2018).

## Materials and Methods

### Participants

Participants were recruited through online advertisements on a national Danish participant recruitment page (www.forsoegsperson.dk) and through an advertisement in a local newspaper. Sixty participants in total were included, 20 in each of the three age groups: 18-26 years old (younger), 39-50 years old (middle) and 69-79 years old (older). Inclusion happened in parallel for all three groups. Exclusion criteria were: MR contraindications, major psychiatric or neurological history, history of drug or alcohol abuse, participation in medical drug testing within six months of the experiment, smoking within three months of the experiment, infectious disease within three weeks of the experiment, morbid obesity, pregnancy, and insufficient understanding of Danish. One participant from the older group was excluded due to an unexpected pathological finding (Table 1).

**Table 1.** Demographics reported as mean ± SD

| Variable | Younger | Middle | Older |
| --- | --- | --- | --- |
| Number of participants (women) | 20 (10) | 20 (10) | 19 (9) |
| Age | 22.6±2.37** | 44.1±3.64** | 72.5±2.78** |
| BMI | 22.6±1.72** | 26.6±4.28* | 26.6±3.45* |
| Years of education | 15.2±2.03 | 16.0±2.82 | 15.1±2.64 |

Note: Age groups that are pairwise significantly different ( $p < 0.05$ ) are marked with \*\* if different from both other groups and \* if different from one other group.

Participants fasted but were allowed to drink water from 22:00h the day before the experiment and until the experiment was over. All participants underwent MR scanning between 9:00h and 11:00h immediately followed by cognitive testing. Participants were informed orally and in writing about the experiment. They provided written consent prior to the experiment. After the experiment, all participants were reimbursed for their time spent participating. The study was approved by the Regional Committee on Health Research Ethics from the Capitol Region in Denmark and was performed in accordance with the declaration of Helsinki (amendment of Fortaleza, 2013).

### MR acquisition

A Phillips 7T whole body MR scanner (Philips, Best, Netherlands) was used in combination with a dual transmit coil and a 32-channel receive head coil (Nova Medical, Wilmington, MA, USA). MR scanning took a maximum of 90 minutes per participant.

A T_1_-weighted magnetization prepared rapid gradient echo (MPRAGE) sequence (slices=380, slice thickness=0.5, TR=8.0 ms, TE=3.2 ms, flip angle=7 degrees, field of view=256×256×190, acquisition matrix=64×64×380) was acquired for anatomical reference and tissue classification.

A semi-localised by adiabatic selective refocusing (sLASER) sequence (Boer et al., 2011; Arteaga de Castro et al., 2013) (TR/TE=3700/32 ms, bandwidth=4 kHz, data points=2048) was used in the medial ACC (20×20×20 mm^3^, 16 acquisitions), left DLPFC (l2×20×20 mm^3^, 32 acquisitions), left hippocampus (30×15×15 mm^3^, 64 acquisitions) and left thalamus(16×12×16 mm^3^, 64 acquisitions) (Figure 1). Variable pulse power and optimized relaxation delay (VAPOR) water suppression was applied (Tkac et al., 2001). A non-water-suppressed spectrum was obtained at the beginning of each sequence. Second order B_0_ shimming was applied using the FASTMAP algorithm (Gruetter and Boesch, 1992; Gruetter, 1993) for all voxels separately using a shim box centred on the voxel and 15 mm longer than the voxel in each direction. Fractions of GM, WM and CSF in each voxel were calculated with voxel-based morphometry using CAT12 (Gaser, C., & Dahnke, 2016) implemented in SPM12 (Statistical Parametric Mapping, Wellcome Department of Cognitive Neurology, London, UK) (Table 2).

**Figure 1:**
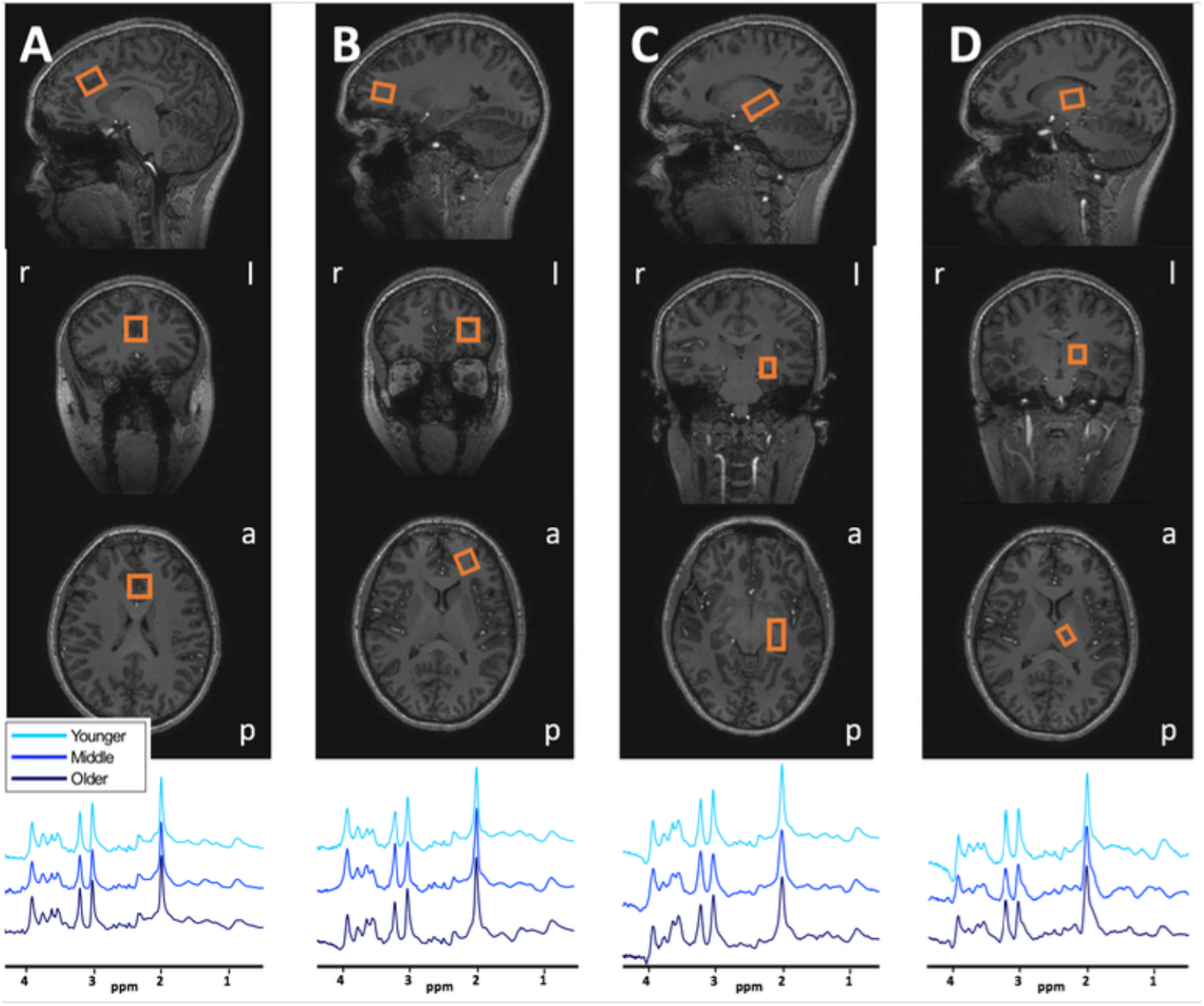
Voxel placements and representative spectra for A) ACC, B) DLPFC, C) Hippocampus D) Thalamus. r: right, l: left, a: anterior, p: posterior.

**Table 2.**
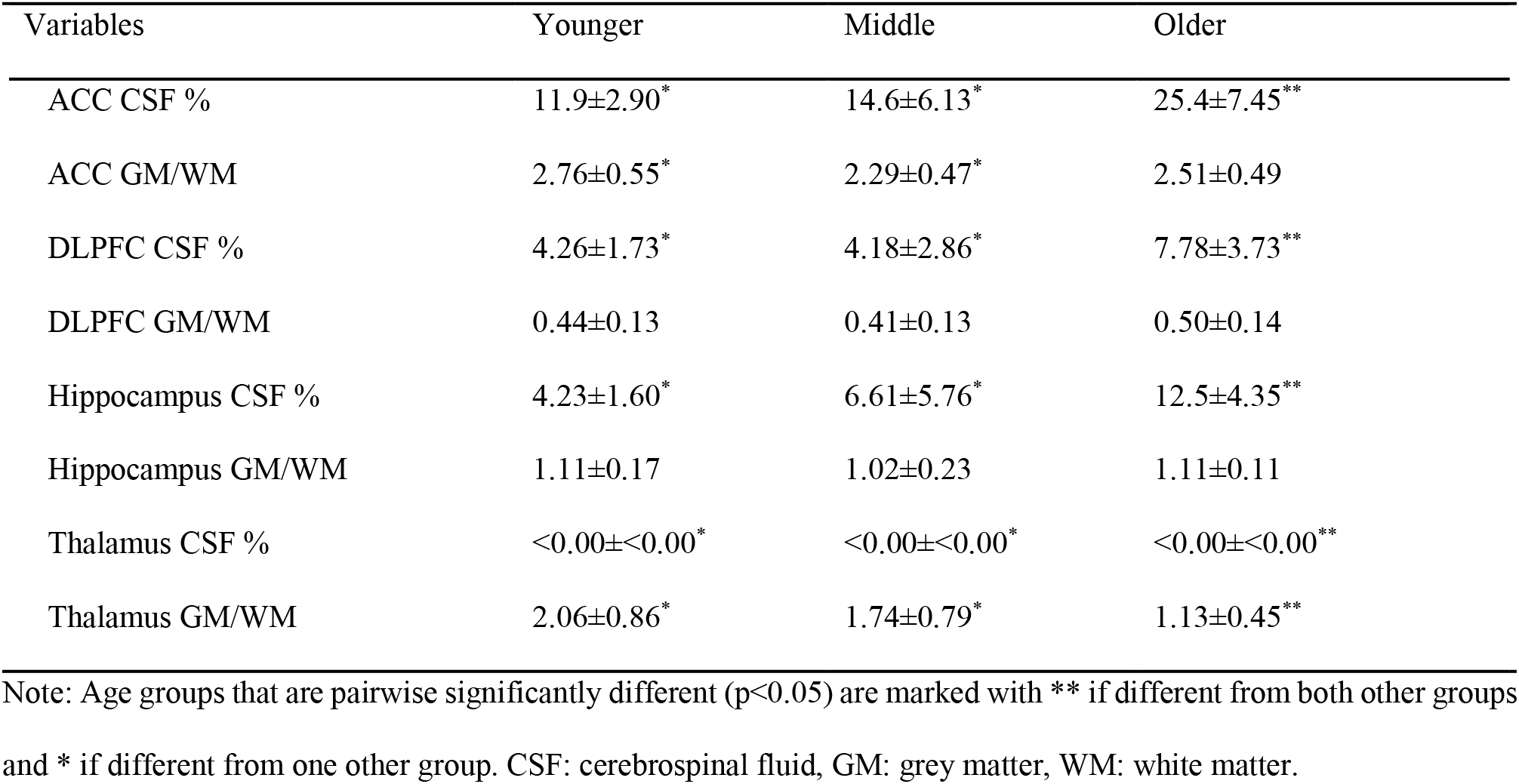
Voxel tissue fractions reported as mean ± SD

| Variables | Younger | Middle | Older |
| --- | --- | --- | --- |
| ACC CSF % | 11.9±2.90* | 14.6±6.13* | 25.4±7.45** |
| ACC GM/WM | 2.76±0.55* | 2.29±0.47* | 2.51±0.49 |
| DLPFC CSF % | 4.26±1.73* | 4.18±2.86* | 7.78±3.73** |
| DLPFC GM/WM | 0.44±0.13 | 0.41±0.13 | 0.50±0.14 |
| Hippocampus CSF % | 4.23±1.60* | 6.61±5.76* | 12.5±4.35** |
| Hippocampus GM/WM | 1.11±0.17 | 1.02±0.23 | 1.11±0.11 |
| Thalamus CSF % | <0.00±<0.00* | <0.00±<0.00* | <0.00±<0.00** |
| Thalamus GM/WM | 2.06±0.86* | 1.74±0.79* | 1.13±0.45** |

Note: Age groups that are pairwise significantly different ( $p < 0.05$ ) are marked with \*\* if different from both other groups and \* if different from one other group. CSF: cerebrospinal fluid, GM: grey matter, WM: white matter.

### Spectral fitting and quantification

After visual inspection, 4% of the spectra were excluded from the analyses due to poor quality; four from the younger group (one from ACC, one from DLPFC, one from hippocampus and one from thalamus) and five from the older group (two from DLPFC, one from hippocampus and two from thalamus). The remaining spectra were fitted with LCModel (Provencher, 2001) using a custom basis set with 20 metabolites (alanine, ascorbate, aspartate (Asp), creatine (Cr), GABA, glutamine (Gln), Glu, glycine, glycerophosphocholine (GPC), glutathione (GSH), mIns, lactate (Lac), N-acetylaspartate (NAA), N-acetylaspartylglutamate (NAAG), phosphorylcholine (PCh), phosphocreatine (PCr), phosphorylethanolamine, scyllo-inositol (sIns), serine (Ser) and taurine (Tau)) including a measured macromolecular baseline. Levels of mIns, tCr (Cr+PCr), Cho (GPC+PCh), tNAA (NAA+NAAG) and Glu were used for the main analyses. The spectra were fitted between 0.2 and 4 ppm with a knot spacing of 0.2. Spectral quality measures were calculated for all spectra including Cramér-Rao lower bounds (CRLB), line widths (FWHM) and signal-to-noise ratios (SNR). Spectral quality was overall good (Table 3). Metabolite quantification was performed through water referencing. Metabolite levels were corrected for tissue fractions in the voxel (Gasparovic et al., 2009; Quadrelli et al., 2016) including tissue specific attenuation factors for T_1_ (Rooney et al., 2007) and T_2_ (Bartha et al., 2002) relaxation times by correcting the water concentration in the voxel (WaterConc_corr_) in the following way:

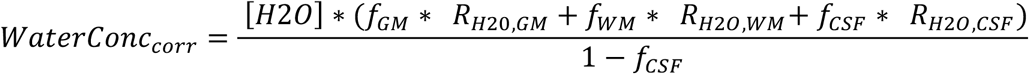

where the water fraction in tissue x, *f*_*X*_, is:

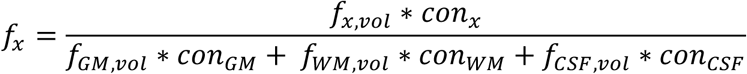

and the tissue specific attenuation factors *R*_*H2O.X*_ is:

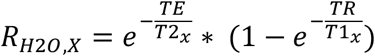

when *[H2O]* is the concentration of pure water, *f*_*x,vol*_ is the fractional volume of tissue x within the voxel, *con*_*x*_ is the water content in tissue x, and *T1*_*x*_ and *T2*_*x*_ are the T_1_ and T_2_ relaxation times of water in tissue x.

**Table 3.**
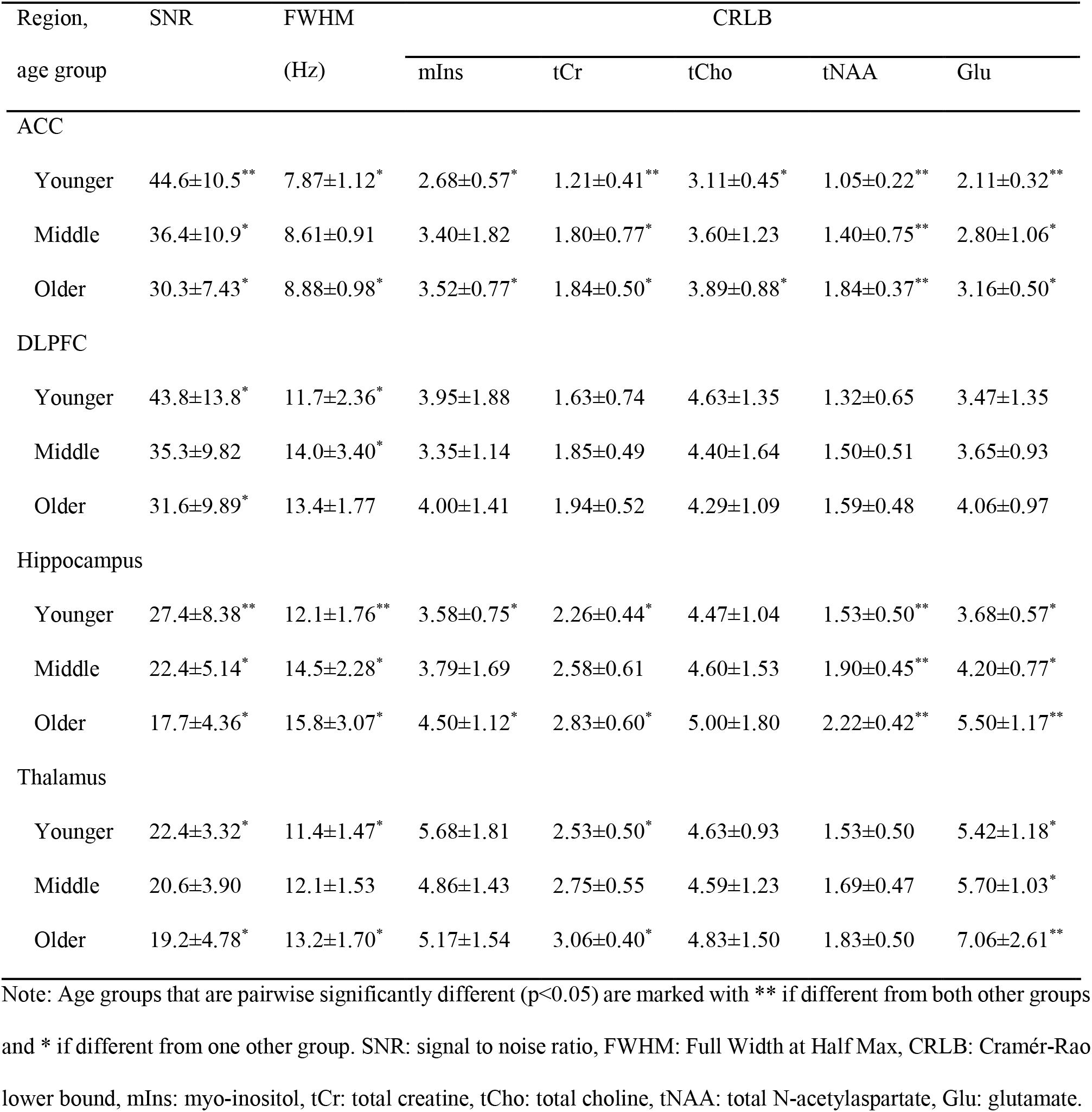
Quality variables for the 1H-MRS measurements reported as mean ± SD

| Region,<br>age group | SNR | FWHM<br>(Hz) | CRLB |  |  |  |  |
| --- | --- | --- | --- | --- | --- | --- | --- |
|  |  |  | mIns | tCr | tCho | tNAA | Glu |
| ACC |  |  |  |  |  |  |  |
| Younger | 44.6±10.5** | 7.87±1.12* | 2.68±0.57* | 1.21±0.41** | 3.11±0.45* | 1.05±0.22** | 2.11±0.32** |
| Middle | 36.4±10.9* | 8.61±0.91 | 3.40±1.82 | 1.80±0.77* | 3.60±1.23 | 1.40±0.75** | 2.80±1.06* |
| Older | 30.3±7.43* | 8.88±0.98* | 3.52±0.77* | 1.84±0.50* | 3.89±0.88* | 1.84±0.37** | 3.16±0.50* |
| DLPFC |  |  |  |  |  |  |  |
| Younger | 43.8±13.8* | 11.7±2.36* | 3.95±1.88 | 1.63±0.74 | 4.63±1.35 | 1.32±0.65 | 3.47±1.35 |
| Middle | 35.3±9.82 | 14.0±3.40* | 3.35±1.14 | 1.85±0.49 | 4.40±1.64 | 1.50±0.51 | 3.65±0.93 |
| Older | 31.6±9.89* | 13.4±1.77 | 4.00±1.41 | 1.94±0.52 | 4.29±1.09 | 1.59±0.48 | 4.06±0.97 |
| Hippocampus |  |  |  |  |  |  |  |
| Younger | 27.4±8.38** | 12.1±1.76** | 3.58±0.75* | 2.26±0.44* | 4.47±1.04 | 1.53±0.50** | 3.68±0.57* |
| Middle | 22.4±5.14* | 14.5±2.28* | 3.79±1.69 | 2.58±0.61 | 4.60±1.53 | 1.90±0.45** | 4.20±0.77* |
| Older | 17.7±4.36* | 15.8±3.07* | 4.50±1.12* | 2.83±0.60* | 5.00±1.80 | 2.22±0.42** | 5.50±1.17** |
| Thalamus |  |  |  |  |  |  |  |
| Younger | 22.4±3.32* | 11.4±1.47* | 5.68±1.81 | 2.53±0.50* | 4.63±0.93 | 1.53±0.50 | 5.42±1.18* |
| Middle | 20.6±3.90 | 12.1±1.53 | 4.86±1.43 | 2.75±0.55 | 4.59±1.23 | 1.69±0.47 | 5.70±1.03* |
| Older | 19.2±4.78* | 13.2±1.70* | 5.17±1.54 | 3.06±0.40* | 4.83±1.50 | 1.83±0.50 | 7.06±2.61** |

Note: Age groups that are pairwise significantly different (p<0.05) are marked with \*\* if different from both other groups and \* if different from one other group. SNR: signal to noise ratio, FWHM: Full Width at Half Max, CRLB: Cramér-Rao lower bound, mIns: myo-inositol, tCr: total creatine, tCho: total choline, tNAA: total N-acetylaspartate, Glu: glutamate.

### Cognitive tests

Cognitive testing was performed on a tablet using a custom-composed Cambridge Neuropsychological Test Automated Battery (CANTAB) (Cambridge Cognition Ltd, Cambridge, United Kingdom) (Sahakian and Owen, 1992). For the present study, two tasks targeting vsWM were included; the paired associates learning (PAL) and the spatial working memory (SWM) task. Both tasks have been shown to be reliable and sensitive to age-related cognitive decline (Robbins et al., 1994; Gonçalves et al., 2016).

### PAL

In PAL, boxes were displayed in a circle. The boxes were opened and closed one by one in a randomised order. Some boxes had a pattern hidden inside. Next, the patterns were displayed one by one in the middle of the screen and the participant had to click the box where the same pattern had previously been displayed. If the correct box for one or more patterns was not chosen, the task was repeated. The number of times a participant selected the wrong box was used as outcome score. Participants had to place all patterns correctly in maximally four attempts to reach the next level with a higher number of patterns. The levels were four, six or eight patterns. If a participant did not reach all levels, the score was adjusted for the levels not reached.

### SWM

In SWM, boxes were displayed in an asymmetrical pattern. The participant had to find a hidden token in as few clicks as possible by opening the boxes one by one. When the token was found, a new token was hidden in one of the other boxes. The outcome score was the number of times a participant selected a box in which a token had previously been found. When tokens had been found in all boxes, the next level was reached. The levels were four, six or eight boxes.

The PAL and SWM scores were correlated (rho=0.478, p<0.001). They were, therefore, z-scored and summed into a vsWM composite score. The vsWM score was inverted so that a higher score represented better performance.

### Experimental Design and Statistical Analysis

To minimise data loss, spectra that were excluded based on poor quality were not considered missing values. Instead, for each metabolite the group mean in that region was imputed. Outliers were defined as metabolite levels more than three SD from the group mean in a given brain region. Outlier detection led to exclusion of one hippocampal tCr and one thalamus mIns value, both from participants in the middle group.

SPSS 25 (Statistical Package for the Social Sciences, Chicago, IL, US) was used for statistical analyses. Threshold of significance was set to p<0.05, after correction for multiple comparisons where applicable. One-way ANOVA testing for the effect of age-group followed by post-hoc pairwise comparisons was applied to compare continuous demographic variables, spectral quality measures, tissue distributions and scores from cognitive testing.

### Overall age-related metabolite differences

A multivariate ANCOVA (MANCOVA) using Pillai’s trace with mIns, tCr, tCho, tNAA and Glu levels as dependent variables was performed to test the primary hypothesis that metabolite levels overall differed across age groups and that there was an interaction between age group and brain region. GM/WM ratio for each voxel was added as covariates.

### Age-related metabolite differences

Next, to investigate the brain region specific differences across age groups, a MANCOVA using Pillai’s trace for each brain region separately was performed, including mIns, tCr, tCho, tNAA and Glu levels as dependent variables. GM/WM ratio in each voxel was added as covariate. Significant main effects were further qualified by *post-hoc* ANCOVA and Bonferroni corrected pairwise comparisons.

### Metabolite correlations with cognition

The secondary hypothesis that age-related differences in metabolites were associated with differences in vsWM was investigated with Bonferroni corrected partial Pearson’s correlations with the GM/WM ratio as covariate. Correlations were tested between vsWM score and metabolites that differed across age groups within a brain region. The tests were one-tailed, testing for a negative correlation between vsWM score and mIns tCr and tCho levels and a positive correlation between vsWM score and tNAA and Glu levels.

### Exploratory metabolite analysis

Several metabolites not included in the main hypotheses were also acquired with ^1^H-MRS. Exploratory statistical analyses of age-related differences for these metabolites were performed per brain region with ANCOVA controlling for GM/WM ratio followed by pairwise comparisons. Metabolites were only included in the analysis if they had at least 50% of the values per group left after removal of values with CRLB > 50 and outliers.

## Results

### Overall age-related metabolite differences

When including all metabolites in all brain regions, there was a significant main effect of age group (F_10,436_=12.49, p<0.001) and a significant age group x brain region interaction (F_30,1105_=1.87, p=0.003). Thus, metabolite levels differ across age groups in a brain region dependent way and each brain region was, therefore, next studied separately.

### Age-related metabolite differences

There was a significant main effect of age group in all separate brain regions, ACC (F_10,104_=4.06, p < 0.001), DLPFC (F_10,104_=4.20, p < 0.001), hippocampus (F_10.102_=6.75, p < 0.001), thalamus (F_10,102_=2.35, p = 0.015). These differences are further qualified with *post-hoc* testing for each metabolite below. See table 4 for metabolite levels and statistics and figure 2 for visualisation. All tests were also done controlling for sex, but this did not significantly affect the results.

**Table 4.**
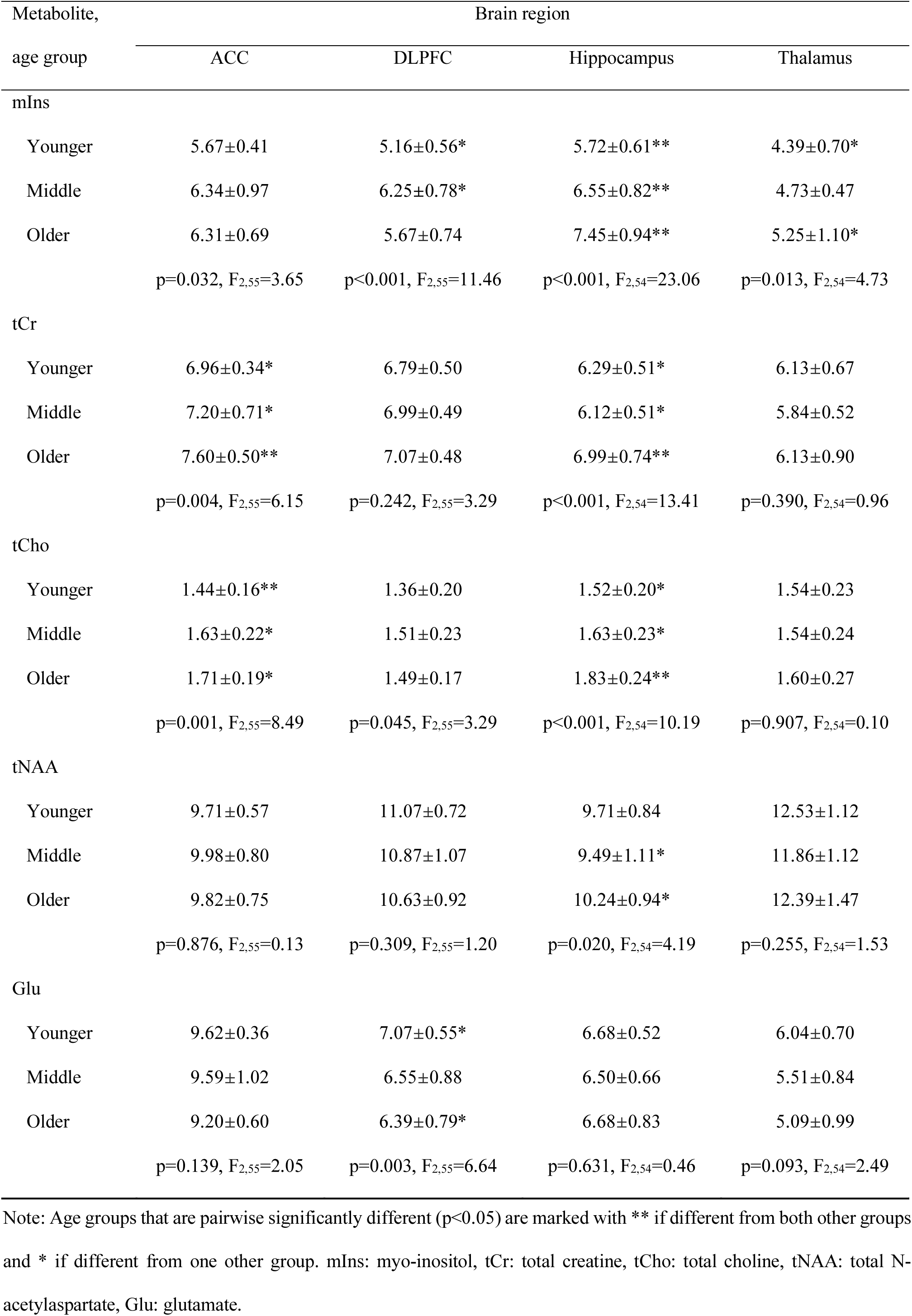
Brain region specific analysis testing for effect of age group. Metabolite levels reported as mean ± SD.

| Metabolite, | Brain region |  |  |  |
| --- | --- | --- | --- | --- |
| age group | ACC | DLPFC | Hippocampus | Thalamus |
| mIns |  |  |  |  |
| Younger | 5.67±0.41 | 5.16±0.56* | 5.72±0.61** | 4.39±0.70* |
| Middle | 6.34±0.97 | 6.25±0.78* | 6.55±0.82** | 4.73±0.47 |
| Older | 6.31±0.69 | 5.67±0.74 | 7.45±0.94** | 5.25±1.10* |
|  | p=0.032, F<sub>2,55</sub>=3.65 | p<0.001, F<sub>2,55</sub>=11.46 | p<0.001, F<sub>2,54</sub>=23.06 | p=0.013, F<sub>2,54</sub>=4.73 |
| tCr |  |  |  |  |
| Younger | 6.96±0.34* | 6.79±0.50 | 6.29±0.51* | 6.13±0.67 |
| Middle | 7.20±0.71* | 6.99±0.49 | 6.12±0.51* | 5.84±0.52 |
| Older | 7.60±0.50** | 7.07±0.48 | 6.99±0.74** | 6.13±0.90 |
|  | p=0.004, F<sub>2,55</sub>=6.15 | p=0.242, F<sub>2,55</sub>=3.29 | p<0.001, F<sub>2,54</sub>=13.41 | p=0.390, F<sub>2,54</sub>=0.96 |
| tCho |  |  |  |  |
| Younger | 1.44±0.16** | 1.36±0.20 | 1.52±0.20* | 1.54±0.23 |
| Middle | 1.63±0.22* | 1.51±0.23 | 1.63±0.23* | 1.54±0.24 |
| Older | 1.71±0.19* | 1.49±0.17 | 1.83±0.24** | 1.60±0.27 |
|  | p=0.001, F<sub>2,55</sub>=8.49 | p=0.045, F<sub>2,55</sub>=3.29 | p<0.001, F<sub>2,54</sub>=10.19 | p=0.907, F<sub>2,54</sub>=0.10 |
| tNAA |  |  |  |  |
| Younger | 9.71±0.57 | 11.07±0.72 | 9.71±0.84 | 12.53±1.12 |
| Middle | 9.98±0.80 | 10.87±1.07 | 9.49±1.11* | 11.86±1.12 |
| Older | 9.82±0.75 | 10.63±0.92 | 10.24±0.94* | 12.39±1.47 |
|  | p=0.876, F<sub>2,55</sub>=0.13 | p=0.309, F<sub>2,55</sub>=1.20 | p=0.020, F<sub>2,54</sub>=4.19 | p=0.255, F<sub>2,54</sub>=1.53 |
| Glu |  |  |  |  |
| Younger | 9.62±0.36 | 7.07±0.55* | 6.68±0.52 | 6.04±0.70 |
| Middle | 9.59±1.02 | 6.55±0.88 | 6.50±0.66 | 5.51±0.84 |
| Older | 9.20±0.60 | 6.39±0.79* | 6.68±0.83 | 5.09±0.99 |
|  | p=0.139, F<sub>2,55</sub>=2.05 | p=0.003, F<sub>2,55</sub>=6.64 | p=0.631, F<sub>2,54</sub>=0.46 | p=0.093, F<sub>2,54</sub>=2.49 |

Note: Age groups that are pairwise significantly different ( $p<0.05$ ) are marked with \*\* if different from both other groups and \* if different from one other group. mIns: myo-inositol, tCr: total creatine, tCho: total choline, tNAA: total N-acetylaspartate, Glu: glutamate.

**Figure 2:**
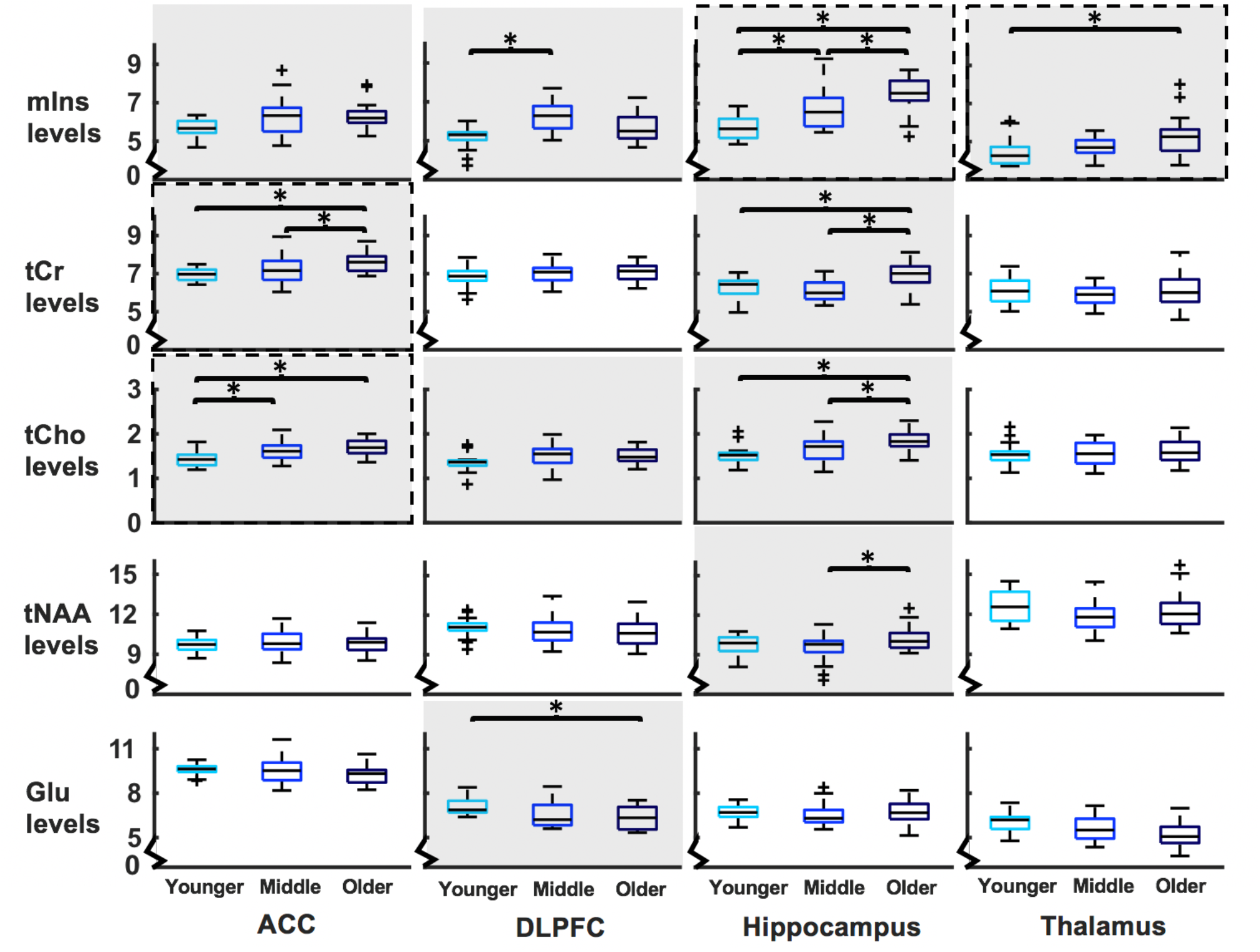
Boxplots of metabolite levels separated by region and age group. Metabolites with a significant main effect of age group at ANCOVA level have a grey background and significant pairwise differences between groups are marked with *. Metabolites that significantly correlate with the visuo-spatial working memory composite score are enclosed in a dashed line. mIns: myo-inositol, tCr: total creatine, tCho: total choline, tNAA: total N-acetylaspartate, Glu: glutamate.

### mIns levels

There was a significant main effect of age group for mIns in ACC (p=0.032, F_2,55_=3.65), DLPFC (p<0.001, F_2,55_=11.46), hippocampus (p<0.001, F_2,54_=23.06) and thalamus (p=0.013, F_2,54_=4.73). In ACC, mIns levels were trendwise higher in the older (mean±SD=6.31±0.69) than the younger group (mean±SD=5.67±0.41, p=0.053) and in the middle (mean±SD=6.34±0.97) than the younger group (p=0.083). In DLPFC, mIns levels were significantly higher in the middle (mean±SD=6.25±0.78) than the younger group (mean±SD=5.16±0.56, p<0.001) and trendwise higher in the older (mean±SD=5.67±0.74) than the younger group (p=0.051). In hippocampus, mIns levels were significantly higher in the older (mean±SD=7.45±0.94) than the younger group (mean±SD=5.72±0.61 p<0.001), in the older than the middle group (mean±SD=6.55±0.82, p=0.002) and in the middle than the younger group (p=0.028). In thalamus, mIns levels were significantly higher in the older (mean±SD=5.25±1.10) than the younger group (mean±SD=4.39±0.70, p=0.010).

### tCr levels

There was a significant main effect of age group for tCr in ACC (p=0.004, F_2,55_=6.15) and hippocampus (p<0.001, F_2,54_=13.41). In ACC, tCr levels were significantly higher in the older (mean±SD=7.60±0.50) than the younger group (mean±SD=6.96±0.34, p=0.005) and the older than the middle group (mean±SD=7.20±0.71, p=0.037). In hippocampus, tCr levels were significantly higher in the older (mean±SD=6.99±0.74) than the younger group (mean±SD=6.29±0.51, p<0.001) and the older than the middle group (mean±SD=6.12±0.51, p<0.001).

### tCho levels

There was a significant main effect of age group for tCho in ACC (p=0.001, F_2,55_=8.49), DLPFC (p=0.045, F_2,55_=3.29) and hippocampus (p<0.001, F_2,54_=10.19). In ACC, tCho levels were significantly higher in the older (mean±SD=1.71±0.19) than the younger group (mean±SD=1.44±0.16, p<0.001) and in the middle (mean±SD=1.63±0.22) than the younger group (p=0.033). In DLPFC, tCho levels were trendwise higher in the older (mean±SD=1.49±0.17) than the younger group (mean±SD=1.36±0.20, p=0.091). In hippocampus, tCho levels were significantly higher in the older (mean±SD=1.83±0.24) than the younger group (mean±SD=1.52±0.20, p<0.001) and the older than the middle group (mean±SD=1.63±0.23, p=0.010).

### tNAA levels

There was a significant main effect of age group for tNAA in hippocampus (p=0.020, F_2,54_=4.19). In the hippocampus, the tNAA levels were significantly higher in the older (mean±SD=10.24±0.94) than the middle group (mean±SD=9.49±1.11, p=0.018).

### Glu

There was a significant main effect of age group for Glu in DLPFC (p=0.003, F_2,55_=6.64). In DLPFC, Glu levels were significantly lower in the older (mean±SD=6.39±0.79) than the younger group (mean±SD=7.07±0.55, p=0.002).

### Metabolite correlations with cognitive ageing

The vsWM, PAL and SWM score all differed across age groups (see table 5 for statistics). vsWM score correlated negatively with mIns in the hippocampus (rho=−0.471, p=0.001) and thalamus (rho=−0.355, p=0.037) and with tCr (rho=−0.364, p=0.027) and tCho (rho=−0.486, p<0.001) in the ACC. See Figure 3 for plots. Before Bonferroni correction, significant correlations were also observed for mIns in ACC (rho=−0.244, p uncorrected=0.032), tCr in hippocampus (rho=−0.263, p uncorrected=0.0424) and Glu in DLPFC (rho=0.269, p uncorrected=0.021).

**Table 5.** Cognitive scores reported as mean ± SD

| Variables | Younger | Middle | Older | Significance test |
| --- | --- | --- | --- | --- |
| vsWM score | 1.30±0.15** | 0.43±0.24** | -1.81±0.39** | $P<0.001$, $F_{2,56}=33.80$ |
| PAL errors | 4.75±0.91* | 12.0±2.15* | 29.6±3.93** | $P<0.001$, $F_{2,56}=24.20$ |
| SWM errors | 3.20±0.98* | 6.65±1.48* | 16.1±2.03** | $P<0.001$, $F_{2,56}=18.41$ |

Note: Age groups that are pairwise significantly different ( $p<0.05$ ) are marked with \*\* if different from both other groups and \* if different from one other group. vsWM: visuo-spatial working memory, PAL: paired associates learning, SWM: spatial working memory

**Figure 3:**
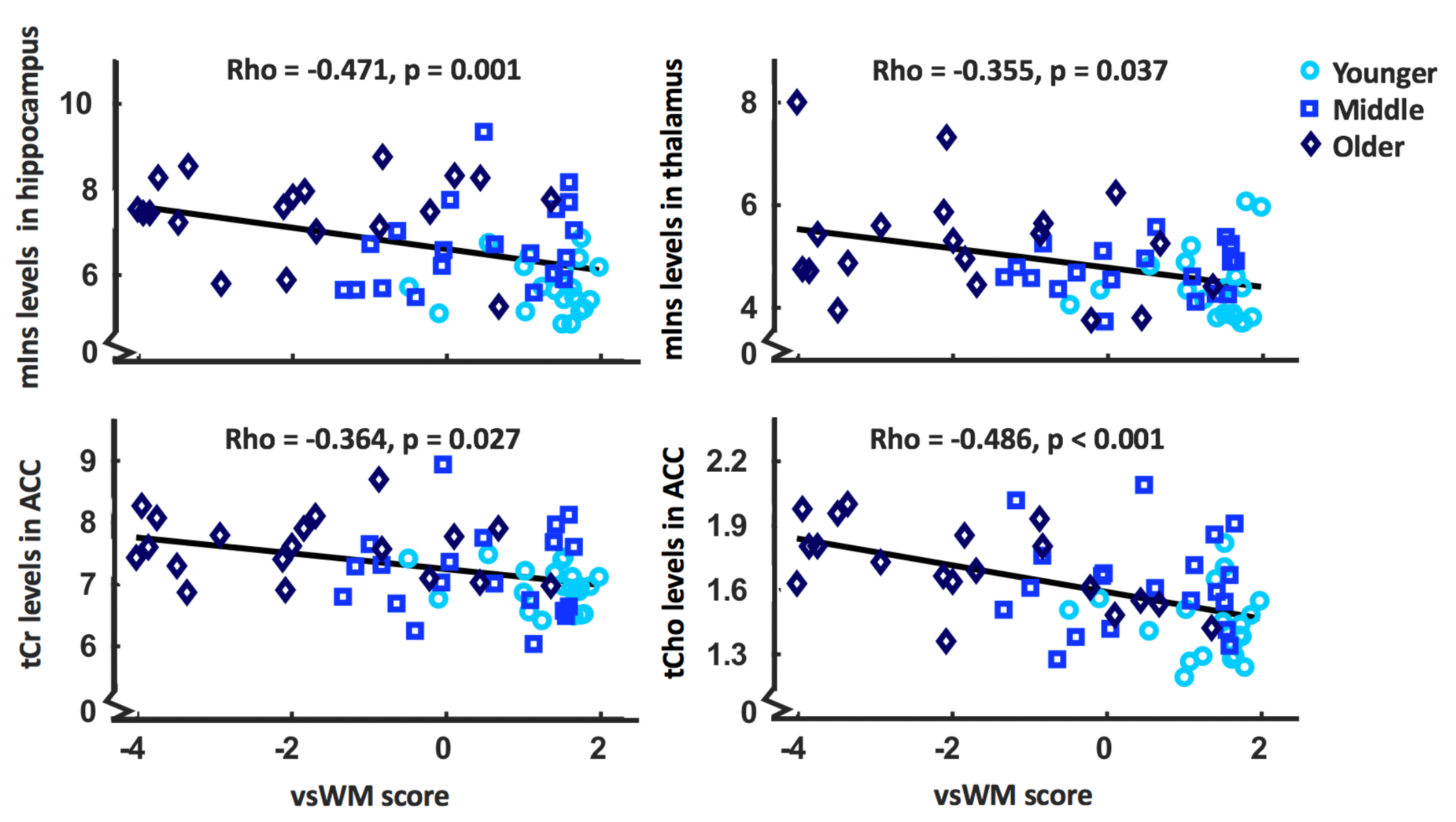
Significant correlations between metabolite levels that differ between age groups and the visuo-spatial working memory score (vsWM). mIns: myo-inositol, tCr: total creatine, tCho: total choline

### Exploratory metabolite analysis

Gln, GSH, NAA, NAAG, sIns and Tau could be included in the analysis within the defined limits in all regions. Additionally, Asp could be quantified in ACC, DLPFC and thalamus, GABA in ACC and thalamus, Lac in DLPFC and Ser in ACC and hippocampus. The exploratory analysis revealed a significant main effect of age group for Gln in hippocampus, Lac in ACC, NAA in DLPFC and thalamus, NAAG in DLPFC and hippocampus, sIns in ACC, DLPFC and hippocampus, Ser in hippocampus and Tau in hippocampus. Except for NAA, levels were higher in older age groups. See table 6 for variable levels and statistics.

**Table 6.**
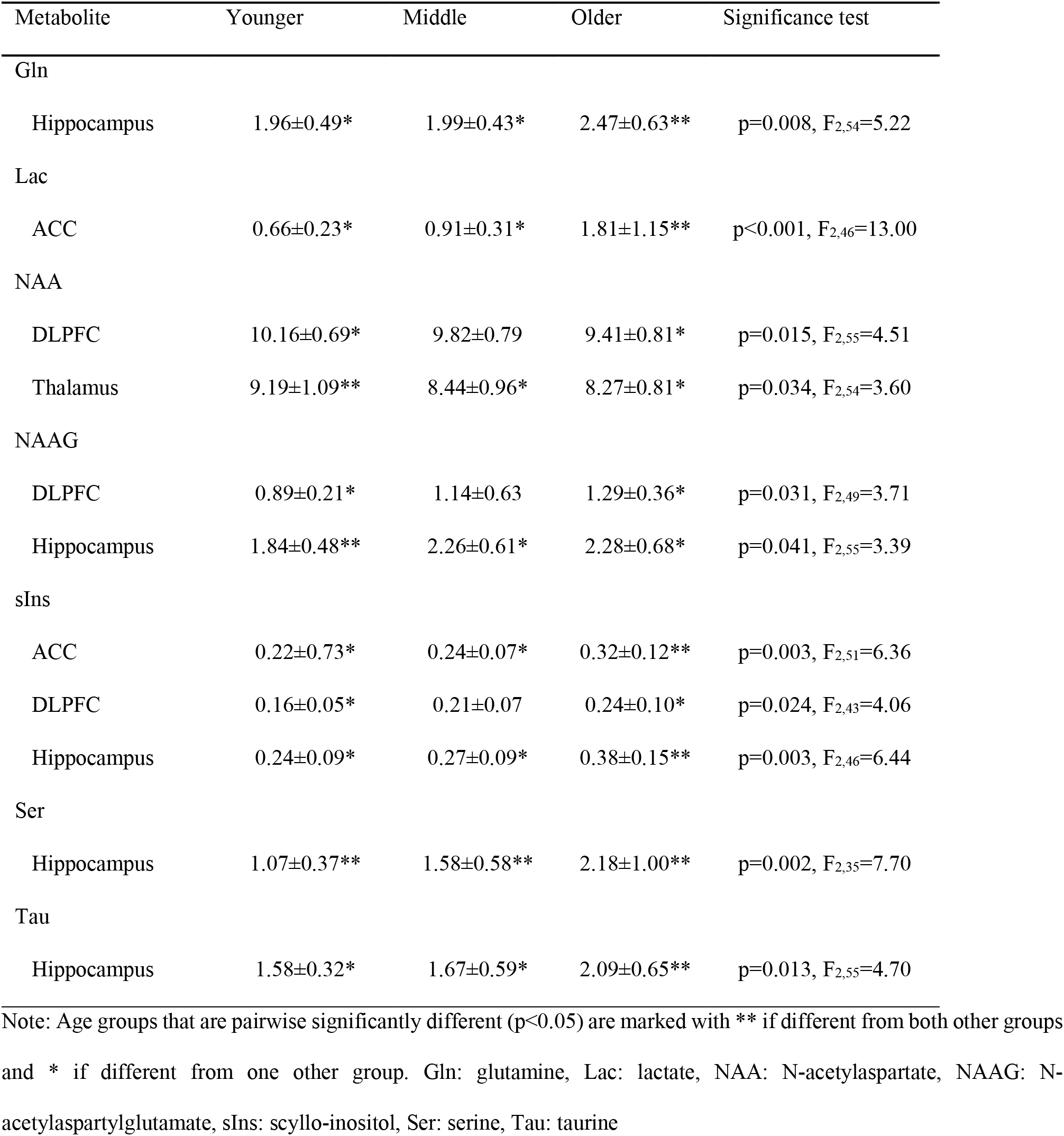
Exploratory analysis testing for effect of age group. Metabolite levels of significant findings reported as mean ± SD.

| Metabolite | Younger | Middle | Older | Significance test |
| --- | --- | --- | --- | --- |
| Gln |  |  |  |  |
| Hippocampus | 1.96±0.49* | 1.99±0.43* | 2.47±0.63** | p=0.008, F<sub>2,54</sub>=5.22 |
| Lac |  |  |  |  |
| ACC | 0.66±0.23* | 0.91±0.31* | 1.81±1.15** | p<0.001, F<sub>2,46</sub>=13.00 |
| NAA |  |  |  |  |
| DLPFC | 10.16±0.69* | 9.82±0.79 | 9.41±0.81* | p=0.015, F<sub>2,55</sub>=4.51 |
| Thalamus | 9.19±1.09** | 8.44±0.96* | 8.27±0.81* | p=0.034, F<sub>2,54</sub>=3.60 |
| NAAG |  |  |  |  |
| DLPFC | 0.89±0.21* | 1.14±0.63 | 1.29±0.36* | p=0.031, F<sub>2,49</sub>=3.71 |
| Hippocampus | 1.84±0.48** | 2.26±0.61* | 2.28±0.68* | p=0.041, F<sub>2,55</sub>=3.39 |
| sIns |  |  |  |  |
| ACC | 0.22±0.73* | 0.24±0.07* | 0.32±0.12** | p=0.003, F<sub>2,51</sub>=6.36 |
| DLPFC | 0.16±0.05* | 0.21±0.07 | 0.24±0.10* | p=0.024, F<sub>2,43</sub>=4.06 |
| Hippocampus | 0.24±0.09* | 0.27±0.09* | 0.38±0.15** | p=0.003, F<sub>2,46</sub>=6.44 |
| Ser |  |  |  |  |
| Hippocampus | 1.07±0.37** | 1.58±0.58** | 2.18±1.00** | p=0.002, F<sub>2,35</sub>=7.70 |
| Tau |  |  |  |  |
| Hippocampus | 1.58±0.32* | 1.67±0.59* | 2.09±0.65** | p=0.013, F<sub>2,55</sub>=4.70 |

Note: Age groups that are pairwise significantly different (p<0.05) are marked with \*\* if different from both other groups and \* if different from one other group. Gln: glutamine, Lac: lactate, NAA: N-acetylaspartate, NAAG: N-acetylaspartylglutamate, sIns: scyllo-inositol, Ser: serine, Tau: taurine

## Discussion

Using 7T ^1^H-MRS, we found differences across age groups on glia-related metabolite levels in ACC, DLPFC, hippocampus and thalamus with the most common difference being higher levels in the older than the younger group. Additionally, glia-related metabolites in ACC, hippocampus and thalamus correlated with cognitive ageing as reflected by vsWM performance. Only few differences were observed in neuronal markers and none of these scaled with vsWM. Together, the results highlight the importance of glial cells in multiple brain regions for cognitive ageing. They also suggest that 7T ^1^H-MRS measurements of glia-related metabolites in key regions of normal brain ageing might be useful in longitudinal studies.

### Glia-related metabolites and cognitive ageing

As hypothesised, regional levels of mIns, tCr and tCho were generally higher in older age groups in the ACC and hippocampus. This corresponds with some previous studies (Chang et al., 1996; Sporn et al., 2019) though other studies observed unaltered levels (Hädel et al., 2013; Ding et al., 2016). Some of the previous negative findings could, however, have arisen from not including participants older than 70 years, low field strengths, the selected scan parameters or insufficient correction for tissue fractions (Cleeland et al., 2019; Sporn et al., 2019). In the DLPFC, levels of mIns, tCr and tCho were not higher in the older than the younger group. As the DLPFC voxel contained more WM than GM, this corresponds with a previous study showing that PFC glia-related metabolite changes during ageing are primarily related to GM (Chang et al., 1996). Overall, the finding of differences in glia-related metabolites across age groups in PFC and hippocampus are consistent with the common notion that these regions are especially sensitive to ageing (Hedden and Gabrieli, 2004).

Thalamus also had higher mIns levels in the older than the younger group. This has not been observed before, however, very few ^1^H-MRS ageing studies have included the thalamus. To our knowledge, this is the first ^1^H-MRS study of normal ageing in thalamus at 7 T and the first to include an age group above 70 years. Accordingly, other MR techniques suggest that the thalamus may be a central node in ageing and could, thus, be of special interest in future ageing studies (Sullivan et al., 2004; Goldstone et al., 2018). Overall, the results indicate that levels of glia-related metabolites are elevated during ageing in PFC, hippocampus and thalamus.

Glia-related metabolite levels and vsWM were correlated for tCho and tCr in ACC and mIns in hippocampus and thalamus. This study is the first to show these correlations during normal ageing, however, only few studies of metabolites and cognitive ageing exist (for review see: (Cleeland et al., 2019)). In PFC, the studies have mainly focused on WM (Ross et al., 2005, 2006; Kochunov et al., 2010). In hippocampus, an association between tNAA/tCr and vsWM has been observed, however, the ratio complicates interpretation (Driscoll et al., 2003). None of the studies included thalamus, however, our results correspond to findings from other types of MR studies suggesting that neurobiological ageing effects on thalamus could be a critical part of cognitive ageing (Grieve et al., 2007; Goldstone et al., 2018). Our study is, thus, the first to pinpoint a specific *in vivo* connection between cognitive ageing of vsWM and elevated levels of glia-related ^1^H-MRS metabolites in ACC, hippocampus and thalamus.

Higher regional levels of mIns, tCr and tCho indicates glia proliferation or gliosis (Glanville et al., 1989; Urenjak et al., 1993). This is consistent with stereological findings that the number of glial cells increase with age in the frontal and temporal cortex (Terry et al., 1987). With age, glial cells, including astrocytes and microglia, change phenotype into a more pro-inflammatory state (Perry et al., 2007; Cohen and Torres, 2019). This increase in neuroinflammation may negatively impact cognitive function (Sartori et al., 2012). As elevations in mIns, tCho and tCr are associated with neuroinflammation, this could be the underlying mechanism for the relationship between age group, metabolites and cognitive ageing (Chang et al., 2013). Additionally, could ageing specifically impact the synthesis and break-down of mIns, tCr and tCho. This would imply that ageing is associated with imbalances in cell signalling, energy homeostasis and membrane turnover respectively (Chang et al., 1996, 2013; Rae, 2014). Overall, these results add to the literature suggesting a central role for glial cells in normal cognitive ageing potentially via neuroinflammation.

### Neuron-related metabolites

Regional levels of tNAA and Glu were not lower in older age groups, except for Glu levels in DLPFC. This corresponds to some studies (Harada et al., 2001; Chang et al., 2009; Haga et al., 2009; Reyngoudt et al., 2012), whereas other studies report lower levels with age (Brooks et al., 2001; Schubert et al., 2004; Ding et al., 2016). Studies reporting lower levels have, however, often quantified the metabolite by calculating the ratio to tCr which could just as well reflect higher tCr levels (Haga et al., 2009). Lower levels of Glu but not tNAA in DLPFC could suggest that neuronal function is altered in this region but not neuronal health. That tNAA remains unaltered in PFC during normal ageing is consistent with stereological observations that frontal neuronal density does not decline with age (Terry et al., 1987). In hippocampus, tNAA levels were even higher rather than lower between the middle and the older group. When previously observed, higher tNAA levels with older age were proposed to arise from the hippocampal neurons’ retained ability for growth in later life and might, thus, represent a beneficial effect (Lie et al., 2004; Bettio et al., 2017; Sporn et al., 2019). On the contrary, lower tNAA levels have been associated with age-related neurodegenerative diseases (Kantarci et al., 2013). Inadvertent recruitment of participants in pre-clinical stages of disease in studies of normal ageing could, thus, have resulted in an apparent tNAA decrease with age. Lower tNAA might, thus, be a trait of pathological rather than normal ageing (Harada et al., 2001; Murray et al., 2014; Wang et al., 2015).

In our exploratory analysis, we used the increased spectral resolution of 7 T ^1^H-MRS to distinguish between NAA and NAAG. Our exploratory analysis indicated that NAA levels were lower in older age groups in DLPFC and thalamus, which was not the case for tNAA. In hippocampus, the results suggest that the higher tNAA levels in the older than middle group could arise from higher NAAG rather than higher NAA. NAAG is a derivative of NAA which might be involved in neuron to glia signalling (Baslow, 2000). It has previously been observed to be associated with ageing in parietal and occipital cortex (Marjańska et al., 2017). The results in this study were only exploratory, however, they support that there could be valuable information in separating NAA and NAAG in future ageing studies.

### Other metabolites

Levels of Gln, Lac and sIns were higher in older age groups in some brain regions. Increased levels with age of these metabolites have been observed before, albeit not necessarily in the same brain regions as used in this study (Sijens et al., 2001; Kaiser et al., 2005; Hädel et al., 2013). Together with the effect on tCr, this could indicate age related differences in energy metabolism (Rae, 2014). The amino acids Tau and Ser also had higher levels in older age groups which, to our knowledge, has not been observed before. This analysis was, however, exploratory and complicated by low metabolite concentrations and overlapping ^1^H-MRS signals (Kaiser et al., 2005). Thus, more studies are needed to clarify the association between these metabolites and ageing.

### Limitations

The study is cross-sectional rather than longitudinal, thus, claims of causality and temporal dynamics, i.e., true decline in metabolites or cognitive performance, cannot be made. Participant recruitment was based on participants responding to an advertisement which might have caused a recruitment bias. Further, only one cognitive domain was used to exemplify cognitive ageing and correlations might thus be specific for vsWM function rather than relating to cognitive ageing in general. Regarding the use of ^1^H-MRS, age and tissue type may affect the T_2_ relaxation time constants of metabolites differently, however, these attenuation factors are unknown for 7 T (Kirov et al., 2008). In relation to tissue type, the setup of the ^1^H-MRS voxel allowed us to control for tissue type across participants but not for distinguishing whether metabolite differences related specifically to one tissue type. Furthermore, the quality of the ^1^H-MRS spectra were generally better for the younger than the older group. However, the overall data quality for the older group was still good.

### Conclusion

This study provides 7T ^1^H-MRS evidence that in ACC, hippocampus and thalamus, older age groups have higher levels of glia-related metabolites and they scale with vsWM performance. These findings emphasise the role for glial cells across multiple brain regions in normal cognitive ageing. We have, thus, identified a set of 7T ^1^H-MRS biomarkers that may help describe cognitive ageing. Understanding normal cognitive ageing is a vital step towards identifying the distinguishing features of successful and pathological ageing.

## Acknowledgment

This project was funded by The Danish Agency for Science, Technology and Innovation grant no. 0601-01370B, and The John and Birthe Meyer Foundation. Hartwig R. Siebner holds a 5-year professorship in precision medicine at the Faculty of Health Sciences and Medicine, University of Copenhagen which is sponsored by the Lundbeck Foundation (Grant Nr. R186-2015-2138).

## Notes

Conflict of interest: Hartwig R. Siebner has received honoraria as speaker from Sanofi Genzyme, Denmark and Novartis, Denmark, as consultant from Sanofi Genzyme, Denmark and as senior editor (NeuroImage) from Elsevier Publishers, Amsterdam, The Netherlands. He has received royalties as book editor from Springer Publishers, Stuttgart, Germany. All other authors declare no conflicts of interest.

